# Tissue- and temperature-dependent expression, enzyme activity, and RNAi knockdown of *Catalase* in a freeze-tolerant insect

**DOI:** 10.1101/2025.02.12.637938

**Authors:** Sarah E. Rokosh, Victoria E. Adams, Robyn Walter, Grace E. Kaiser, Amber L. Gough, Jantina Toxopeus

## Abstract

Organisms that overwinter in temperate climates may experience freezing and freezing-induced oxidative stress during winter. While many insect species can survive freezing, molecular tools such as RNA interference (RNAi) or CRISPR have not been used to understand the physiological mechanisms underlying freeze tolerance. The spring field cricket *Gryllus veletis* can survive freezing following a 6-week fall-like acclimation. We used RNAi of an antioxidant enzyme in *G. veletis* to test the hypothesis that minimizing oxidative stress is important for freeze tolerance. In fat body tissue, *Catalase* mRNA abundance and enzyme activity increased during the acclimation that induces freeze tolerance. Other tissues such as midgut and Malpighian tubules had more stable or lower *Catalase* expression and activity during acclimation. In unacclimated (freeze-intolerant) crickets, RNA interference (RNAi) effectively knocked down production of the *Catalase* mRNA and protein in fat body and midgut, but not Malpighian tubules. In acclimated (freeze-tolerant) crickets, RNAi efficacy was temperature-dependent, functioning well at warm (c. 22°C) but not cool (15°C or lower) temperatures. This highlights a challenge of using RNAi in cold-acclimated organisms, as they may need to be warmed up for RNAi to work, potentially affecting their stress physiology. Knockdown of *Catalase* via RNAi in acclimated crickets also had no effect on the ability of the crickets to survive a mild freeze treatment, suggesting that Catalase may not be necessary for freeze tolerance. Our study is the first to demonstrate that RNAi is possible in a freeze-tolerant insect, but further research is needed to examine whether other genes and antioxidant molecules are important in freeze tolerance of *G. veletis*.

**Highlights:**

- *Catalase* expression and activity are elevated in freeze-tolerant cricket fat body
- RNAi knocks down *Catalase* in fat body and midgut at a warm temperature (22°C)
- RNAi is not effective at a cool temperature (15°C) that preserves freeze tolerance
- *Catalase* knockdown has no impact on survival of a mild freeze treatment
- The role of antioxidants in freeze tolerance warrants further study

**Graphical abstract:** 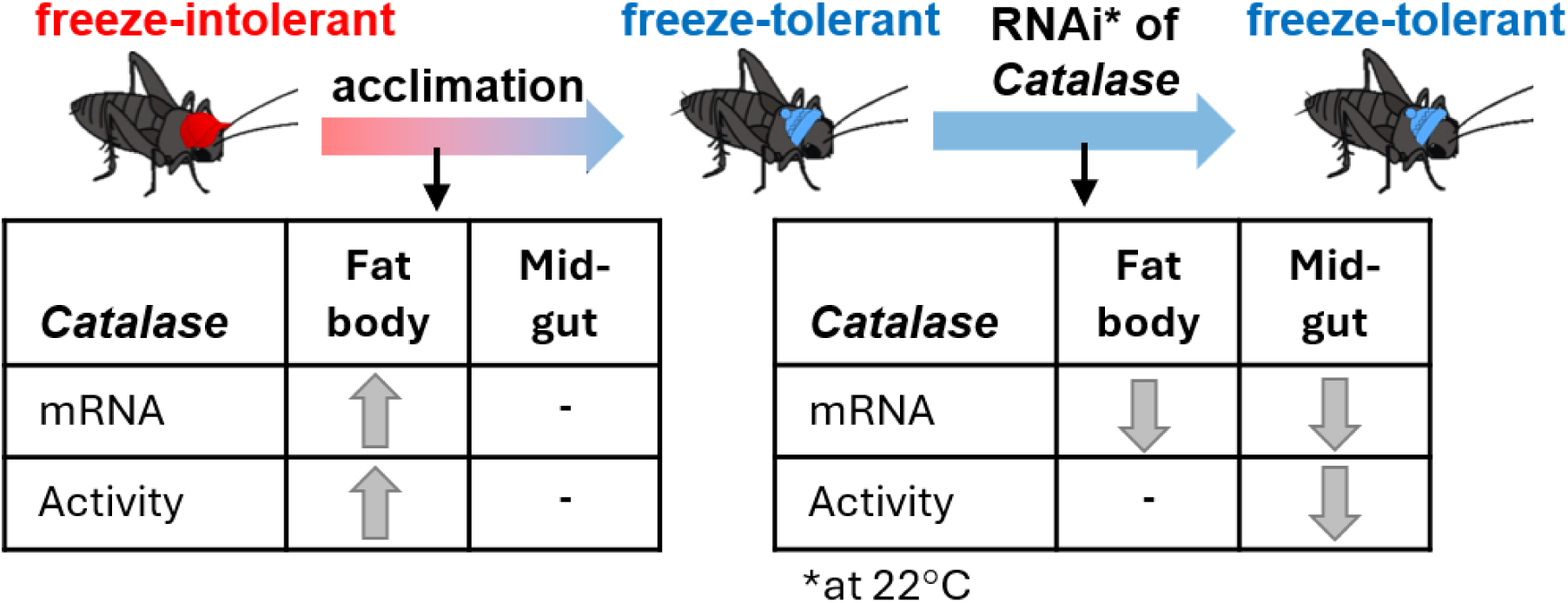

## Introduction

Freezing is lethal to most organisms, but many insects are freeze-tolerant and can survive internal ice formation. Freezing and thawing are hypothesized to cause multiple challenges, including those associated with low temperatures, ice itself, changes in osmolarity, metabolic limitations, and oxidative damage (Rozsypal, 2022; Storey and Storey, 2010, 2013; Teets et al., 2023; Toxopeus and Sinclair, 2018). With this complex system of potentially-interacting stressors, many of the physiological mechanisms underlying freeze tolerance are not well understood (Toxopeus and Sinclair, 2018). Molecular tools such as RNA interference (RNAi) or CRISPR offer an opportunity to study these mechanisms, but have not been used to examine freeze tolerance, in part because few freeze-tolerant species are amenable to laboratory rearing and manipulation (Lebenzon and Toxopeus, 2024; Toxopeus and Sinclair, 2018). However, the spring field cricket *Gryllus veletis* (Orthoptera: Gryllidae) (Alexander & Bigelow) is a freeze-tolerant laboratory model organism (Adams et al., 2025; Smith et al., 2021; Toxopeus et al., 2019a, 2019c, 2019b; van Oirschot and Toxopeus, 2024) that is likely amenable to RNAi (Toxopeus, 2018), similar to other crickets (Meyering-Vos et al., 2006; Miyawaki et al., 2004). *G. veletis* become freeze-tolerant as fifth instar nymphs when exposed to fall or fall-like conditions for at least 6 weeks (Adams et al., 2025; Toxopeus et al., 2019c). Previous transcriptomic work on the fat body tissue during this acclimation identified many upregulated genes that may support freeze tolerance, including antioxidants that can mitigate oxidative damage (Toxopeus et al., 2019a). Our study focuses on whether mechanisms known to be involved in oxidative stress tolerance can also support freeze tolerance.

Oxidative stress arises when production of reactive oxygen species (ROS) exceeds cellular capacity to neutralize ROS via antioxidant mechanisms (Birben et al., 2012; Sokolova, 2018). There are multiple types of ROS, all of which can be derived from molecular oxygen (O_2_) via normal cellular processes (Hancock, 2009; Shields et al., 2021). For example, hydrogen peroxide (H_2_O_2_) is produced via catabolism of fatty acids in organelles called peroxisomes, where it is then neutralized to water (H_2_O) and O_2_ in a reaction catalyzed by the antioxidant enzyme Catalase (Shields et al., 2021). Superoxide (O_2−_) is often produced via the mitochondrial electron transport system (Lebenzon et al., 2023), where is it usually converted to the less reactive H_2_O_2_ via the antioxidant enzyme Superoxide Dismutase (Blier et al., 2014; Shields et al., 2021). While O_2−_ is more reactive than H_2_O_2_, the latter is more mobile because it can diffuse across cellular and organelle membranes (Birben et al., 2012). The H_2_O_2_ produced by Superoxide Dismutase must be further processed to H_2_O via other antioxidant enzymes to fully neutralize this ROS system (Birben et al., 2012; Shields et al., 2021). In addition to a suite of antioxidant enzymes, antioxidant molecules such as glutathione are important for neutralizing ROS (Birben et al., 2012; Shields et al., 2021; Storey and Storey, 2010). If not neutralized, ROS react with cellular macromolecules (proteins, lipids, nucleic acids), causing damage that may be irreparable such as protein carbonylation and denaturation, lipid peroxidation and compromised membrane integrity, and double-stranded breaks in DNA (Shields et al., 2021).

Freezing and thawing are hypothesized to cause oxidative stress in multiple ways. Low temperatures can impair the function of enzymes (Somero et al., 2017), allowing increased rates of ROS accumulation due to reduced antioxidant enzyme activity (Lalouette et al., 2011; Rojas and Leopold, 1996). Low temperatures and freezing can also damage or impair the electron transport system of mitochondria (Cormier et al., 2022; Lebenzon et al., 2023; McMullen and Storey, 2008), potentially increasing ROS production. When ice forms, increased intracellular concentrations of ions such as Fe^2+^ may facilitate ROS production via the Fenton reaction (Storey and Storey, 2010). Frozen insects typically accumulate metabolites associated with anaerobic metabolism (Michaud et al., 2008; Storey and Storey, 1985, 1981), suggesting they are oxygen-limited in this state. ROS production may increase during thawing, as aerobic metabolism involving oxygen increases (Storey and Storey, 2010), especially if mitochondria were damaged during freezing (e.g., Štětina et al., 2020). While a few studies have shown that oxidative damage accumulates after freezing (especially repeated freezing; Doelling et al., 2014), it is not clear whether this is a general phenomenon, or whether oxidative stress tolerance is important for freeze tolerance.

Following the framework of Lebenzon and Toxopeus (2024), we developed and used an RNAi protocol in *G. veletis* to test the hypothesis that oxidative stress tolerance is important for freeze tolerance. We initially selected *Catalase* as a potential target for RNAi because it is upregulated at the mRNA level in the fat body of crickets during fall-like acclimation (Toxopeus et al., 2019a) and is a known antioxidant enzyme. Prior to RNAi, we tested whether *Catalase* expression and enzyme activity were increased during acclimation in multiple tissues (fat body, Malpighian tubules, midgut) that exhibit *Catalase* expression or activity in other insects (Felton and Duffey, 1991; Rajarapu et al., 2011; Zhang et al., 2016).We then designed and delivered long dsRNA targeting *Catalase* for knockdown into *G. veletis*, tested the efficacy of knockdown in the same three tissue types (fat body, Malpighian tubules, midgut) at two different temperatures, and then used the most effective protocol to knock down *Catalase* in acclimated (freeze-tolerant) crickets. We predicted that knocking down *Catalase* would reduce the oxidative stress tolerance of *G. veletis*, and therefore decrease their ability to survive freezing.

## Methods

### Insect rearing and acclimation

All experiments were conducted on male fifth instar nymphs of *G. veletis.* Crickets were reared under summer-like conditions using previously described methods (Toxopeus et al., 2019c) in the Animal Care Facility at St. Francis Xavier University in Antigonish, Nova Scotia, Canada from a population that was originally collected in Lethbridge, Alberta, Canada. To induce freeze tolerance, male fifth instar nymphs were haphazardly selected and exposed to a fall-like acclimation with decreasing temperature and photoperiod over six weeks (Fig. 1A) as previously described (Adams et al., 2025; van Oirschot and Toxopeus, 2024). MIR154 incubators (PHCbi; Wood Dale, IL, USA) were used for acclimation and post-acclimation temperature exposures, unless described otherwise.

**Figure 1.**
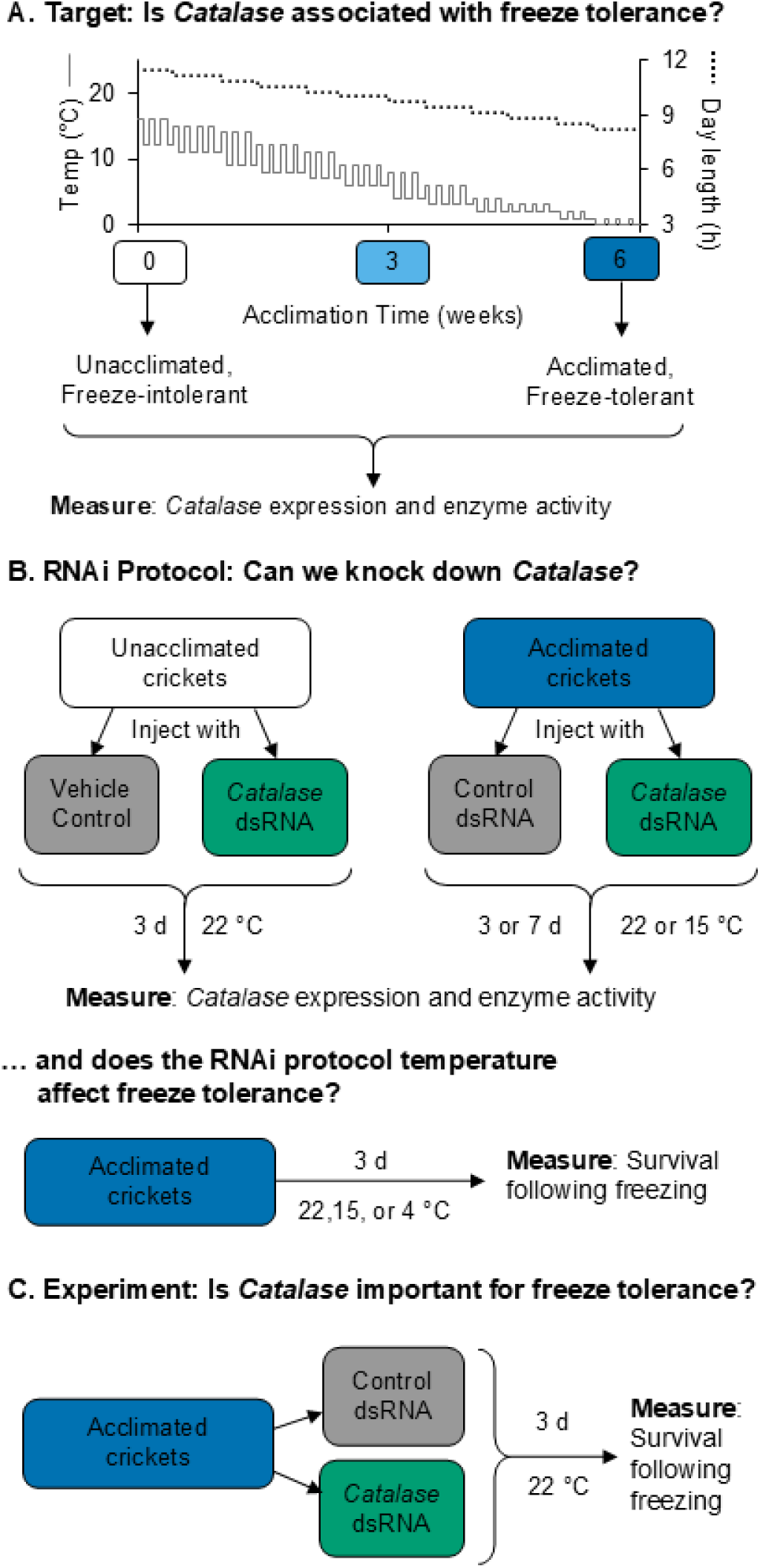
Experimental design to (A) select target a gene (*Catalase*) for RNA interference (RNAi), (B) optimize the RNAi protocol, and (C) test the effect of *Catalase* knock down on freeze tolerance of *Gryllus veletis*. **(A)** The temperatures (daily fluctuations) and day lengths for the 6-week acclimation that induces freeze tolerance are shown. **(B)** A preliminary RNAi pilot was conducted in unacclimated (freeze-intolerant) crickets before testing a wider range of protocols in acclimated (freeze-tolerant crickets). **(C)** The most effective RNAi protocol was used to test whether *Catalase* knockdown affected freeze tolerance. Colour-coding of groups here is consistent with colour-coding in Results figures.

### Experimental designs and sample preparation

To determine changes in *Catalase* expression and enzyme activity during acclimation (Fig. 1A), we selected crickets after 0 weeks (unacclimated), 3 weeks, and 6 weeks of the fall-like acclimation (Adams et al., 2025; van Oirschot and Toxopeus, 2024). We dissected three types of tissues (fat body, Malpighian tubules, midgut) from *G. veletis* either at room temperature (c. 22°C; for unacclimated crickets) or on ice (after 3 or 6 weeks acclimation). Each biological replicate contained tissues pooled from three crickets. All samples were flash-frozen in liquid nitrogen after pooling, and stored at -80°C until processing for RT-qPCR or enzyme assays.

To assess the effect of knocking down *Catalase* on freeze tolerance, we first tested the efficacy of RNAi in the crickets under different conditions (Fig. 1B). Unacclimated and acclimated nymphs were injected with dsRNA targeting *Catalase*, a vehicle control, or a control dsRNA construct targeting *green fluorescent protein* (*GFP*) – a gene that does not exist in these crickets (Lebenzon and Toxopeus, 2024; Toxopeus, 2018). The relative abundance of *Catalase* mRNA and activity of Catalase enzyme were determined in tissues dissected from crickets 3 or 7 d post-injection. Post-injection crickets were either maintained at a warm temperature (room temperature; c. 22°C) or moderate low temperature (15°C). For testing RNAi efficacy, each biological replicate was tissue from a single cricket 3 or 7 d post-injection. All tissues were flash-frozen in liquid nitrogen and stored at -80°C until processing for RT-qPCR or enzyme activity assays. RNAi protocols that resulted in successful knockdown at the mRNA and protein level were then repeated in new cohorts of acclimated crickets to test whether *Catalase* knockdown affected their ability to survive a mild freeze treatment (1.5 h at -8°C; Fig. 1C).

To test whether the temperature exposures in the RNAi protocol affected freeze tolerance, we also exposed acclimated crickets that were never injected with dsRNA but were exposed to different post-acclimation conditions (Fig. 1B). Following the fall-like acclimation, groups of nymphs were exposed to 4°C (similar to the end of acclimation), 15°C (similar to the beginning of acclimation), or c. 22°C (room) temperature for 3 d. We then tested their ability to survive a moderate freeze treatment (1.5 h at -8°C).

### Measuring relative mRNA abundance with RT-qPCR

To determine changes in *Catalase* mRNA abundance during acclimation and following RNAi, we extracted RNA from the samples described in the experimental designs, and conducted RT-qPCR. Each sample was homogenized in 300 μL TRIzol (ThermoFisher; Mississauga, ON, Canada), and RNA was extracted according to the manufacturer’s protocol, with a final elution of RNA in 50 μL of nuclease-free water (Adams et al., 2025). Quality and quantity of RNA was then determined by measuring absorbance at 230 nm, 260 nm, and 280 nm in a NanoDropOne microvolume spectrophotometer (ThermoFisher). We used DNAse I (ThermoFisher) to degrade any contaminating genomic DNA, and then performed cDNA synthesis on 100 – 200 ng RNA using the iScript Reverse Transcriptase Supermix (BioRad; Mississauga, ON, Canada) according to manufacturers’ instructions. We diluted cDNA samples 1/10 prior to use in RT-qPCR.

We ordered our RT-qPCR primers from ThermoFisher. We designed the *Catalase* primers (forward: 5’-GCCCGACAATGTTATCCACC-3’; reverse: 5’-TGCACCAAACTACTTCCCCA- 3’) to be complementary to a *G. veletis Catalase* transcript (NCBI accession GGSD01082898.1) using the Primer3 tool (https://primer3.ut.ee/). *Elongation factor 1α* (*EF1α*) was chosen as a reference gene (forward: 5’-CGCTCAATATGGTTGTTGGA-3’; reverse: 5’- CCATCGAAAGATTTGATGTGG-3’), based on previous success in *G. veletis* (Toxopeus, 2018). PCR amplicons were generated with each primer pair and *G. veletis* cDNA using DreamTaq DNA polymerase (ThermoFisher) according to manufacturer’s instructions (Lemay et al., 2024; McIntyre et al., 2023). To verify that our RT-qPCR primers were specific for our targets, we sent the amplicons for Sanger sequencing at the Toronto Centre for Applied Genomics (Toronto, ON, Canada).

We ran RT-qPCR in triplicate for each RNA sample described in the experimental designs, using each of the *Ef1α* and *Catalase* primers under conditions that supported primer efficiency between 90 and 110% (Adams et al., 2025). RT-qPCR was performed according to the manufacturer’s protocol (SsoAdvanced SYBR Green Supermix; BioRad) in 10 µL reaction mixes with 2 µL diluted cDNA, 400 nM primers and the following parameters: initial denaturation at 94°C for 5 min; 40 cycles of 94°C for 15 s, 63°C (*Ef1α*) or 60°C (*Catalase*) for 15 s, 72°C for 20 s; followed by melt curve determination via ramping from 65°C to 95°C in increments of 0.5°C. We used the 2^-ΔΔC^_T_ method (Livak and Schmittgen, 2001) to determine whether relative abundance of *Catalase* mRNA changed during acclimation (compared to control crickets with 0 weeks acclimation) or following RNAi (compared to crickets injected with vehicle/control dsRNA).

### Measuring Catalase enzyme activity

To determine changes in Catalase enzyme activity during acclimation and following RNAi, we generated cell-free protein extracts from the samples described in the experimental designs, and conducted enzyme activity assays (Orr and Sohal, 1992; Pichaud et al., 2010). Each sample was homogenized in 100 mM potassium phosphate buffer (17.4 mg/mL K_2_HPO_4_, 13.6 mg/mL KH_2_PO_4_ in deionized water; pH 6.9) on ice using a plastic pestle. The homogenized tissues were centrifuged at 3000 × *g* for 5 min at 4°C, supernatants were transferred into new tubes on ice and centrifuged at 5000 × *g* for 30 min. The resulting cell-free supernatant was further diluted 1/20 in potassium phosphate buffer for use in BCA and enzyme activity assays. We used the Pierce BCA Protein Kit (ThermoFisher) to spectrophotometrically determine total protein concentration (mg/mL) in the extracts following the manufacturer’s protocol and using a SpectraMaxR iD3 spectrophotometer (Molecular Devices; San Jose, CA, USA).

We used a Catalase enzyme activity assay previously described (Orr and Sohal, 1992; Pichaud et al., 2010). Briefly, we added 5 μL of diluted protein extract and 125 μL of 60 mM H_2_O_2_ (freshly prepared from 10% w/v H_2_O_2_; Sigma-Alrich; St. Louis, MO, USA) in triplicate into half-area ultraviolet (UV) microplates. The samples were mixed for 30 s at a low intensity inside the spectrophotometer, and absorbance of H_2_O_2_ (240 nM) was then measured every 16 s at 22°C for 5 min. We calculated the Catalase enzyme activity using the following formula:

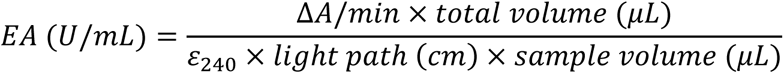

where *EA* is the Catalase enzyme activity in activity units (U, µmol H_2_O_2_ consumed per min) per mL, *ΔA/min* is the absolute value of the linear slope of H_2_O_2_ absorbance at 240 nm over time (min^-1^), *total volume* is the volume of sample in the UV plate wells (130 µL), ε_240_ is the molar absorption of H_2_O_2_ at 240 nm (43.6 mLcm^-1^µmol^-1^), *light path* is the distance travelled by the light through the sample wells in (0.662 cm), and *sample volume* is the volume of sample in the wells (5 µL). The enzyme activity of each sample was calculated as an average of the technical triplicates, and was then standardized to the total protein concentration in the sample (U/mg protein). In RNAi experiments, the enzyme activity relative to controls (crickets injected with control dsRNA constructs) was calculated to make protein and mRNA results more comparable.

### RNA interference protocol

To conduct RNAi, we synthesized dsRNA constructs, injected those constructs into nymphs, and waited 3 to 7 d to allow time for knockdown of *Catalase*. To synthesize dsRNA, we used the T7 MEGAscript RNAi Kit (ThermoFisher), which requires a dsDNA template that includes a T7 promoter region for dsRNA synthesis via T7 reverse transcriptase. To synthesize the dsDNA template for *Catalase*, we used DreamTaq DNA Polymerase to conduct PCR according to manufacturer’s instructions with 2 ng/μL of template (*G. veletis* cDNA) and 1 μM each of forward primer (5’-<u>TAATACGACTCACTATAGGAG</u>CTGGTAATTTGACACT-3’) and reverse primer (5’-<u>TAATACGACTCACTATAGGGAGA</u>CCCAGATTATAGCAT-3’), designed using the same software as above and with a T7 promoter sequence (underlined) incorporated. To synthesize dsDNA template for *GFP*, we conducted PCR using 2 ng/μL template (CRISPR Universal Negative Control 1, a commercially-available vector; Sigma) and 1 μM each of forward primer (5’- <u>TAATACGACTCACTATAGGGAG</u>CTGGAGATCAAGTTCATC-3’) and reverse primer (5’- <u>TAATACGACTCACTATAGGGAG</u>TTAGCCTCCAGGCCGTTG-3’), designed to be complementary to the *GFP* sequence in the commercially-available vector. We used the following reaction conditions: an initial denaturation of 95°C for 5 min; 5 cycles of 95°C for 30 s, 60°C for 30 s, and 72°C for 1 min; 35 cycles of 95°C for 30 s, 65°C for 30 s, 72°C for 1 min; and a final extension at 72°C for 5 min (Toxopeus et al., 2014). The resulting dsDNA templates were used at 0.1 μg/μL to synthesize dsRNA in a single reaction at 37°C for 4 h using the T7 Megascript RNAi Kit, followed by degradation of dsRNA and any ssRNA via DNase and RNase at 37°C for 1 h according to manufacturer’s instructions. dsRNA constructs were diluted to 0.3 μg/μL in nuclease-free water prior, and then 5 μL of diluted *Catalase* dsRNA or control *GFP* dsRNA construct was injected into cricket hemolymph under the pronotum using a 10 μL gastight Hamilton syringe and a 30-gauge disposable needle (Toxopeus, 2018). In early pilot experiments with unacclimated crickets, a vehicle control (5 μL nuclease-free water) was used instead of *GFP* dsRNA (see Fig. 1 and Results figure captions). Crickets were kept in individual mesh-covered clear plastic containers with food and water and shelter (egg carton) until either dissected for validation of *Catalase* knockdown or used in experiments testing cricket survival following freezing.

### Assessing freeze tolerance

We froze crickets as described previously (Adams et al., 2025; van Oirschot and Toxopeus, 2024) under conditions that usually result in >90% survival of fall-acclimated (freeze-tolerant) crickets (Toxopeus et al., 2019c). Briefly, crickets were placed in individual 1.7 mL tubes in an aluminum block cooled by an Arctic A25 recirculating chiller (ThermoFisher) containing 50% (v/v) propylene glycol. 2 μL of silver iodide suspended in distilled water was applied to the dorsal side of each cricket to promote freezing (Adams et al., 2025; Toxopeus et al., 2019c; van Oirschot and Toxopeus, 2024). Crickets were equilibrated at 4°C for 10 min, then cooled to -8°C at -0.25°C/min, held at -8°C for 90 min, then rewarmed to 4°C at 0.25°C/min. We confirmed that the crickets froze (Sinclair et al., 2015) by recording their temperatures once per second using T-type thermocouples (Omega Engineering, Norwalk, CT, USA) interfaced to Picolog v6.2.7 software (Pico Technology, Cambridge, UK) via TC-08 units. At the end of the freeze treatment, crickets were transferred to 15°C for recovery in individual containers as described above. Crickets were classified as alive if they were motile or responded to gentle prodding after 48 h of recovery.

### Statistical analyses

To compare relative *Catalase* mRNA abundance and enzyme activity between crickets acclimated for 0, 3, and 6 weeks, we conducted a one-way ANOVA for each tissue on ΔΔC_T_ values and EA/mg protein values, respectively. If the ANOVA was significant (α = 0.05), we conducted a Tukey’s post-hoc test. To test whether RNAi knocked down *Catalase* mRNA abundance and enzyme activity after 3 d at 22°C in unacclimated crickets, we conducted one-tailed *t*-tests for each tissue on ΔΔC_T_ values and EA/mg protein values, respectively. To test whether the duration (3 or 7 d) and temperature (22°C or 15°C) affected RNAi efficacy in acclimated crickets, we conducted two-way ANOVAs for each temperature and tissue with RNAi treatment (*Catalase* or *GFP* dsRNA) and duration as factors. To assess whether the proportion of freeze-tolerant crickets differed among post-acclimation temperature treatments, or between acclimated crickets injected with *Catalase* or *GFP* dsRNA, we conducted χ^2^ tests.

## Results

### Catalase expression and activity increases in fat body tissue during acclimation

We observed tissue-specific changes in the *Catalase* mRNA abundance and enzyme activity (Fig. 2). In fat body tissue *Catalase* mRNA abundance increased approximately 4-fold mid-way through the fall-like acclimation (Fig. 2A; ANOVA: *F*_2,13_ = 9.702, *P* = 0.003), and enzyme activity increased by approximately 30% between the beginning and end of acclimation (Fig. 2B; ANOVA: *F*_2,15_ = 6.364, *P* = 0.023). Conversely, in midgut tissue both *Catalase* mRNA abundance (Fig. 2A; ANOVA: *F*_2,19_ = 1.749, *P* = 0.201) and enzyme activity (Fig. 2B; ANOVA: *F*_2,12_ = 0.134, *P* = 0.720) remained stable throughout acclimation. In Malpighian tubules *Catalase* mRNA abundance decreased approximately 3-fold mid-acclimation and remained low until the end of acclimation (Fig. 2A; ANOVA: *F*_2,15_ = 6.121, *P* = 0.010), while enzyme activity was unchanged over the same time period (Fig. 2B; ANOVA: *F*_2,13_ = 0.389, *P* = 0.543).

**Figure 2.**
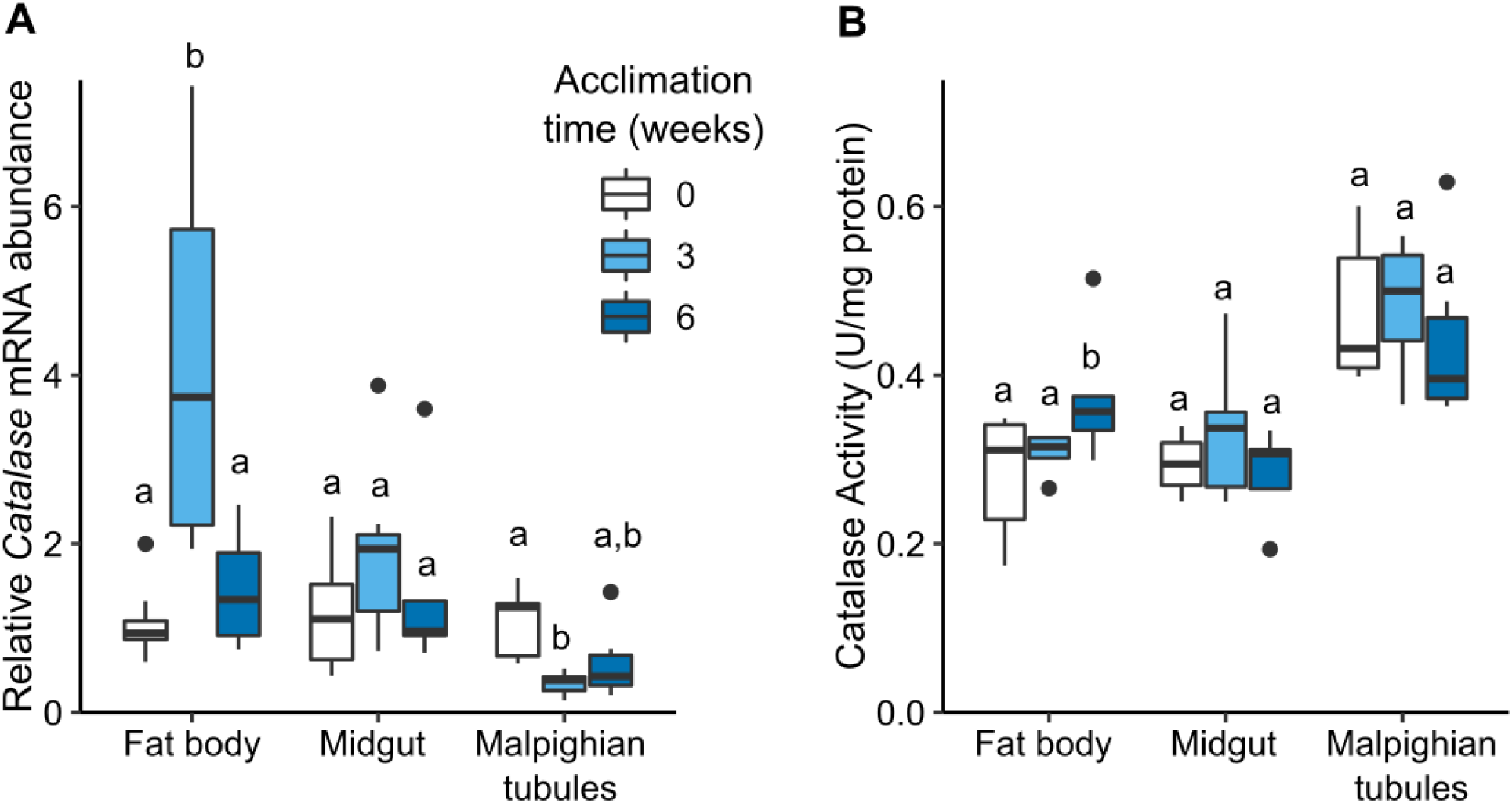
Tissue-specific changes in *Gryllus veletis* (A) *Catalase* mRNA abundance and (B) enzyme activity during fall-like acclimation that induces freeze tolerance. Crickets were either unacclimated (0 weeks), partially acclimated (3 weeks), or fully acclimated (6 weeks). Relative mRNA abundance was determined via RT-qPCR using rpl18 as a reference gene. Enzyme activity was determined using spectrophotometric assays of H2O2 consumption. Each treatment group included 7 – 8 crickets. Within a tissue, different letters indicate statistical differences based on ANOVA with Tukey’s post-hoc test (P < 0.05).

### RNAi is effective in fat body and midgut tissues of unacclimated cricket

We observed tissue-specific efficacy of RNAi targeting *Catalase* at room temperature (c. 22°C) 3 d after unacclimated crickets were injected with dsRNA constructs (Fig. 3). In fat body tissue, RNAi was effective and caused approximately a 75 % decrease in *Catalase* mRNA abundance (Fig. 3A; t-test: *t*_7_ = -4.610, *P* = 0.001), and 45% decrease in enzyme activity (Fig. 3B; t-test: *t*_16_ = 3.748, *P* = 0.001). Similarly, in midgut tissue *Catalase* mRNA abundance was approximately 65 % lower (Fig. 3A; t-test: *t*_13_ = -4.043, *P* < 0.001) and enzyme activity was approximately 55% lower (Fig. 3B; t-test: *t*_17_ = 3.576, *P* = 0.001) in RNAi crickets relative to control crickets. In Malpighian tubules, the *Catalase*-targeting dsRNA construct was effective at the mRNA level (Fig. 3A; t-test: *t*_12_ = -5.838, *P* < 0.001), but ineffective at decreasing enzyme activity (Fig. 3B; t-test: *t*_22_ = 0.831, *P* = 0.208).

**Figure 3.**
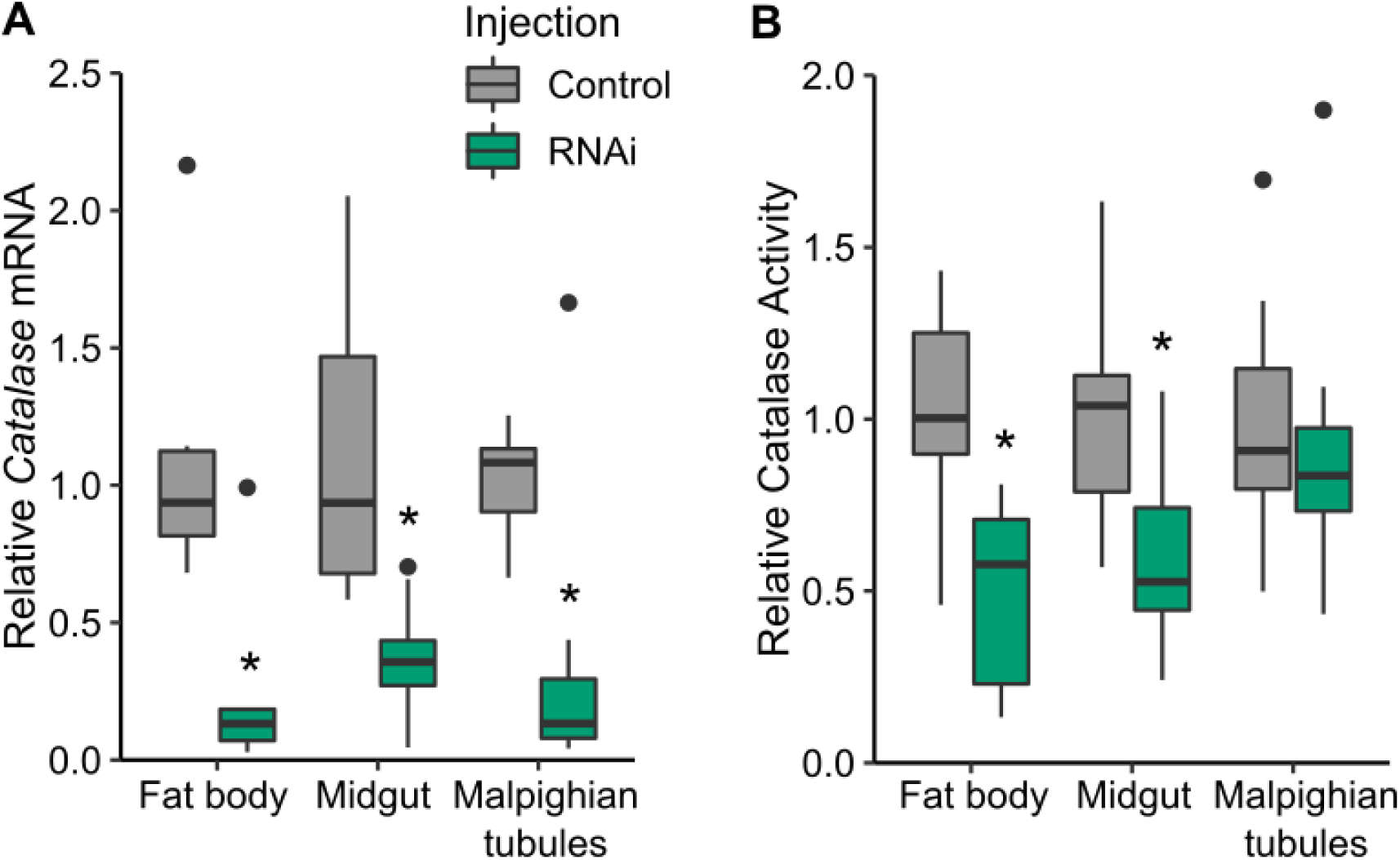
Tissue-specific efficacy of *Catalase* RNA interference (RNAi) in unacclimated *Gryllus veletis* based on (A) *Catalase* mRNA abundance and (B) enzyme activity following injection with dsRNA constructs. Unacclimated crickets were injected with dsRNA targeting *Catalase* (RNAi) or a vehicle control (water), and remained at room temperature (c. 22°C) for 3 d prior to dissection. Relative mRNA abundance was determined via RT-qPCR using *rpl18* as a reference gene. Relative enzyme activity was determined using spectrophotometric assays of H_2_O_2_ consumption. Each treatment group included 7 – 8 crickets. Within a tissue, asterisks indicate statistically-significant knockdown based on a one-tailed Welch’s two sample t-test (*P* < 0.05).

### Acclimated crickets can lose freeze tolerance following short, warm incubations

The freeze tolerance of acclimated crickets was only maintained if crickets were held at low temperature following completion of the 6-week fall-like acclimation (Fig. 4). Freeze tolerance was high (at least 70% survival) if acclimated crickets were maintained at temperatures of 4°C or 15°C for 3 d post-acclimation, but post-freeze survival decreased below 40% if the crickets were exposed to room temperature (c. 22°C) for the same duration (Fig. 4; χ^2^_2,81_ = 8.555, *P* = 0.014). While the crickets used in this experiment were not injected with anything, the results suggested that the temperature used in RNAi protocol with unacclimated crickets (Fig. 2) could potentially impair freeze tolerance of acclimated crickets.

**Figure 4.**
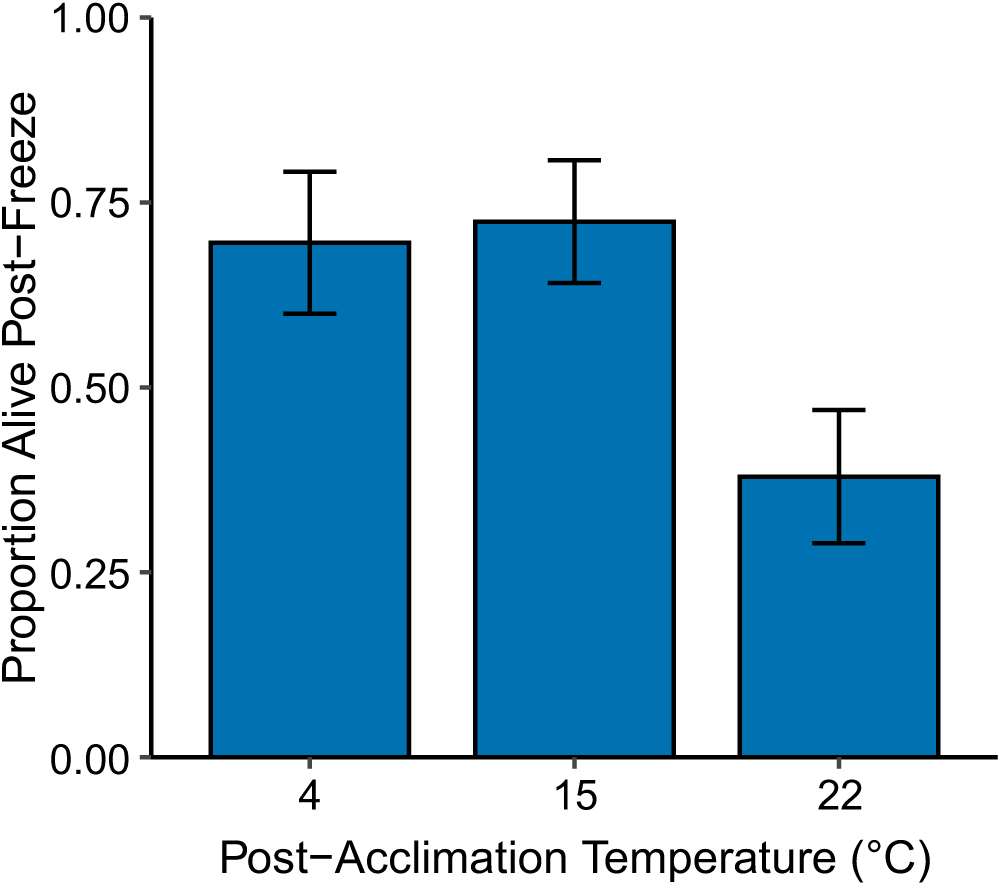
The effect post-acclimation incubation temperature on *Gryllus veletis* freeze tolerance. Acclimated crickets were transferred to 4°C (similar to the end of acclimation), 15°C (similar to the beginning of acclimation), or c. 22°C (room temperature) for 3 d prior to testing freeze tolerance. The proportion of crickets that survived freezing was determined 48 h after exposing them to a 1.5 h freeze treatment at -8°C. Error bars represent the standard error of proportion. Each treatment group included 23 – 30 crickets. Proportion survival differed between temperatures based on a χ^2^ test (*P* < 0.05).

### RNAi is effective in acclimated crickets at warm but not cool temperatures

Similar to unacclimated crickets (Fig. 3), RNAi was effective in acclimated cricket fat body and midgut when these crickets were maintained at a warm temperature (c. 22°C) post-injection (Figs. 5, 6). At these warm temperatures, *Catalase* mRNA in fat body tissue was lower in RNAi than control crickets 3 and 7 d post-injection (Fig. 5A; ANOVA: RNAi treatment *F*_1,28_ = 7.240, *P* = 0.012; Time post-injection *F*_1,28_ = 2.168, *P* = 0.005; Treatment × Time *F*_1,28_ = 1.763, *P* = 0.199). Enzyme activity was statistically unaffected by the RNAi treatment at warm temperatures, although there was a trend toward reduced activity 3 d post-injection with dsRNA construct targeting *Catalase* (Fig. 5B; ANOVA: RNAi treatment *F*_1,22_ = 1.098, *P* = 0.435; Time post-injection *F*_1,22_ = 0.005, *P* = 0.942; Treatment × Time *F*_1,22_ = 0.948, *P* = 0.341). RNAi was successful in midgut tissue of crickets that were maintained at room temperature (warm; c. 22°C) for 3 or 7 d post-injection, with decreased *Catalase* mRNA (Fig. 6A; ANOVA: RNAi treatment *F*_1,260_ = 23.597, *P* < 0.001; Time post-injection *F*_1,26_ = 2.193, *P* = 0.151; Treatment × Time *F*_1,26_ = 2.187, *P* = 0.151) and Catalase enzyme activity (Fig. 6B; ANOVA: RNAi treatment *F*_1,22_ = 28.56, *P* < 0.001; Time post-injection *F*_1,22_ = 18.55, *P* < 0.001; Treatment × Time *F*_1,22_ = 1.930, *P* = 0.179) compared to control crickets.

**Figure 5.**
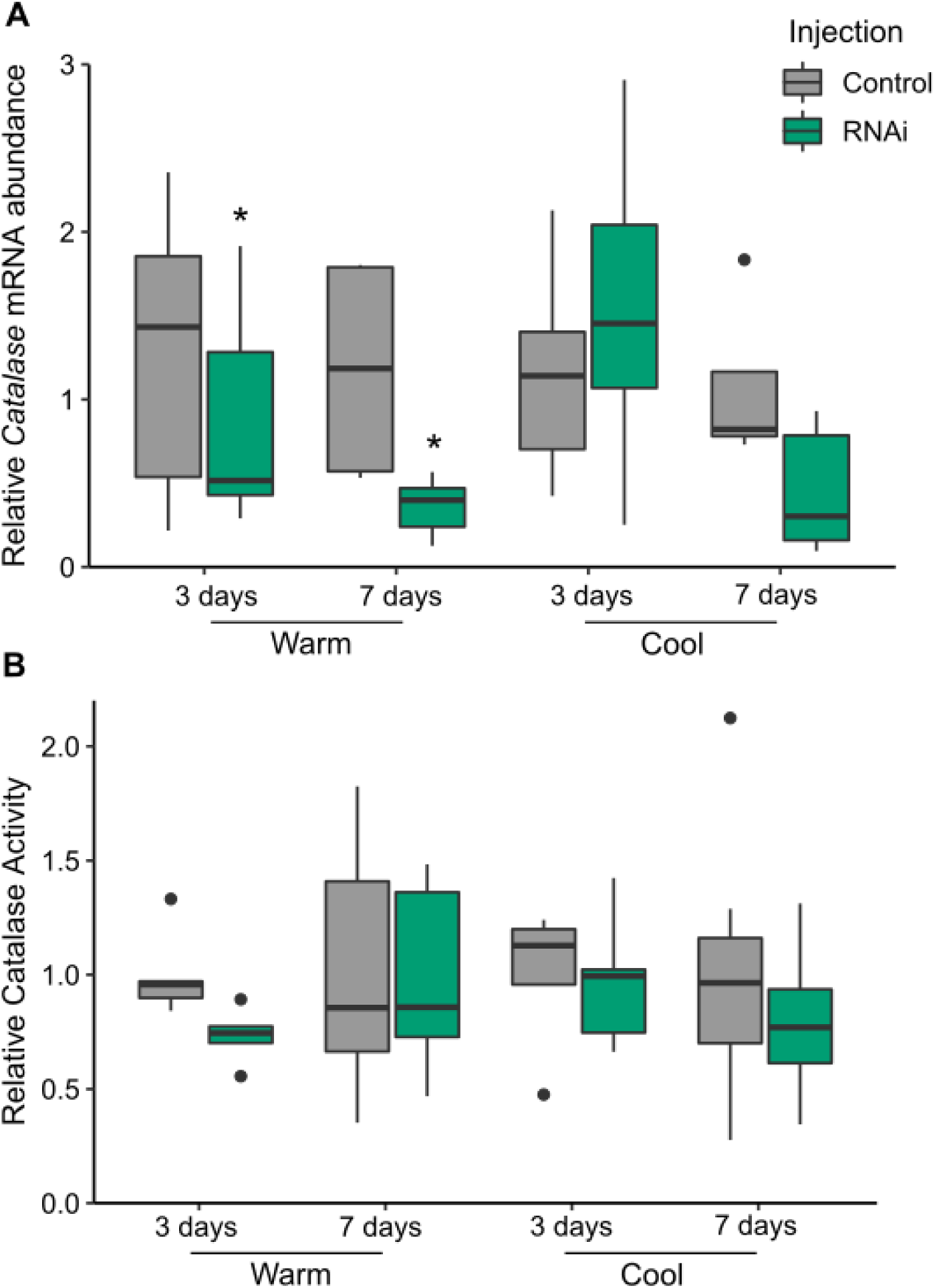
The effect of post-injection temperature and time on efficacy of *Catalase* RNA interference (RNAi) in acclimated *Gryllus veletis* fat body tissue based on (A) *Catalase* mRNA abundance and (B) enzyme activity. Acclimated crickets were injected with dsRNA targeting *Catalase* (RNAi) or a control construct, and remained at a warm (c. 22°C) or cool (15°C) temperature for 3 – 7 days prior to dissection. Relative mRNA abundance was determined via RT-qPCR using *rpl18* as a reference gene. Relative enzyme activity was determined using spectrophotometric assays of H_2_O_2_ consumption. Each treatment group included 6 – 12 crickets. Asterisks indicate a statistically-significant knockdown due to RNAi at the indicated temperature based on a two-way ANOVA (*P* < 0.05).

**Figure 6.**
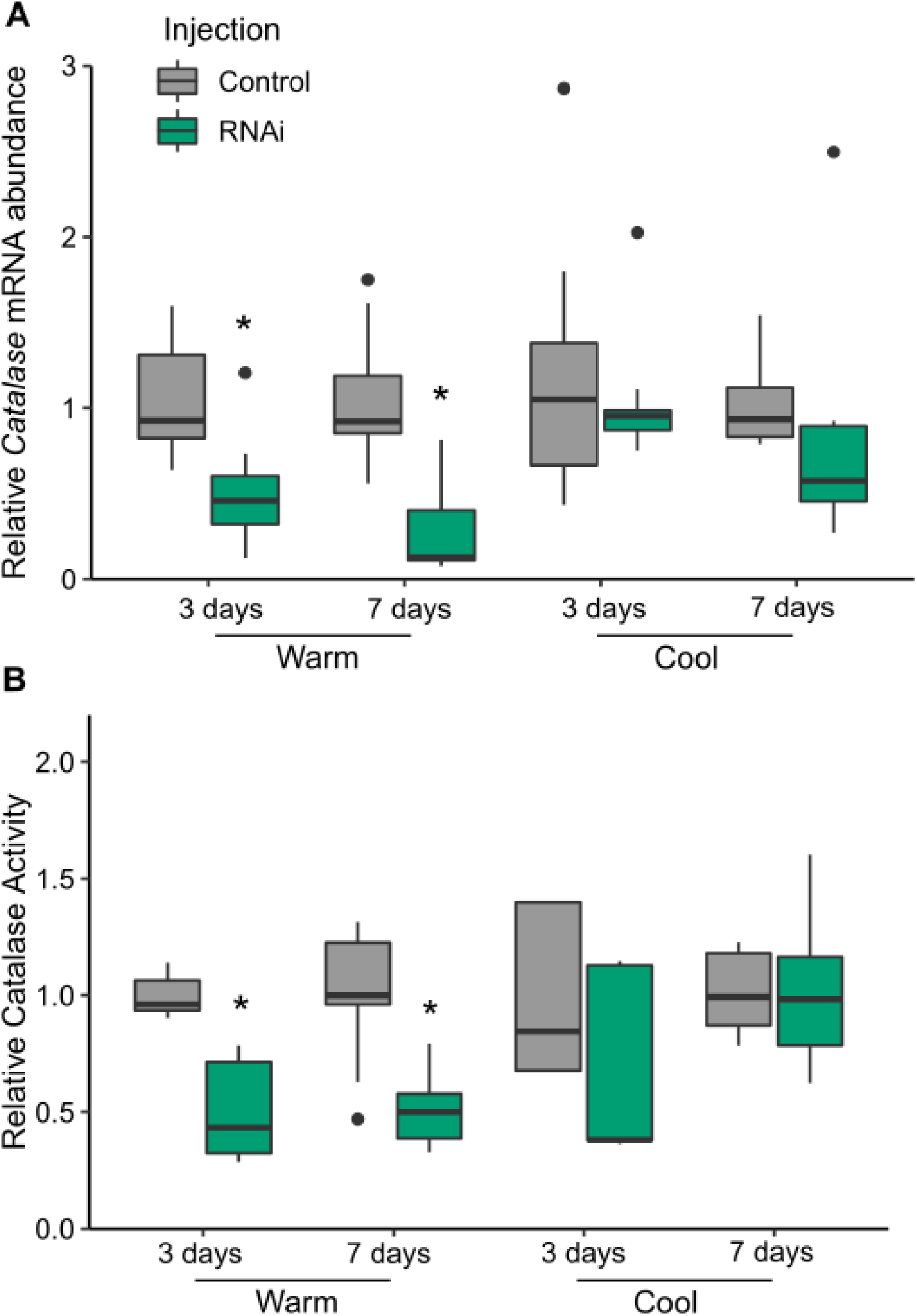
The effect of post-injection temperature and time on efficacy of *Catalase* RNA interference (RNAi) in acclimated *Gryllus veletis* midgut tissue based on (A) *Catalase* mRNA abundance and (B) enzyme activity. Acclimated crickets were injected with dsRNA targeting *Catalase* (RNAi) or a control construct at room temperature, and remained at a warm (c. 22°C) or cool (15°C) temperature for 3 – 7 days prior to dissection. Relative mRNA abundance was determined via RT-qPCR using *rpl18* as a reference gene. Relative enzyme activity was determined using spectrophotometric assays of H_2_O_2_ consumption. Each treatment included 6 – 12 crickets. Asterisks indicate a statistically-significant knockdown due to RNAi at the indicated temperature based on a two-way ANOVA (*P* < 0.05).

However, when RNAi was attempted in acclimated crickets at a cool temperature (15°C) permissive to freeze tolerance (Fig. 4), efficacy was greatly reduced compared to RNAi conducted at a warm temperature (Figs. 5, 6). In fat body, *Catalase* mRNA abundance was not affected by RNAi at cool temperatures 3 or 7 d post-injection (Fig. 5A; ANOVA: RNAi treatment *F*_1,30_ = 1.003, *P* = 0.325; Time post-injection *F*_1,30_ = 9.328, *P* = 0.005; Treatment × Time *F*_1,30_ = 8.016, *P* = 0.008), nor was enzyme activity (Fig. 5B; ANOVA: RNAi treatment *F*_1,25_ = 1.098, *P* = 0.305; Time post-injection *F*_1,25_ = 0.430, *P* = 0.518; Treatment × Time *F*_1,25_ = 0.411, *P* = 0.527). Similarly, *Catalase* knockdown was not effective in midgut tissue after either duration at cool temperatures at the mRNA level (Fig. 6A; ANOVA: RNAi treatment *F*_1,29_ = 1.277, *P* = 0.268; Time post-injection *F*_1,29_ = 1.463, *P* = 0.236; Treatment × Time *F*_1,29_ = 1.296, *P* = 0.264) and the protein level (Fig. 6B; ANOVA: RNAi treatment *F*_1,25_ = 0.392, *P* = 0.567; Time post-injection *F*_1,25_ = 15.800, *P* < 0.001; Treatment × Time *F*_1,25_ = 1.338, *P* = 0.258). There was a trend towards lower *Catalase* mRNA abundance 7 d post-injection in the RNAi group at 15°C in both tissues (Figs. 5B, 6B), suggesting that longer post-injection incubations at cool temperatures have the potential for effective RNAi.

### Catalase RNAi does not affect survival of a mild freeze treatment

Although incubating acclimated crickets at 22°C decreases freeze tolerance (Fig. 4), this was the only condition under which RNAi was effective (Figs. 5, 6), so we used an RNAi protocol at room temperature to test whether Catalase was important for freeze tolerance. When we conducted RNAi targeting *Catalase* in acclimated crickets, we saw no effect of this RNAi treatment on the crickets’ ability to survive freezing for 1.5 h at -8°C relative to the control treatment (Fig. 7; χ^2^_1,43_ = 0.201, *P* = 0.654). Survival of crickets was low (c. 50%) in both injection groups relative to usual survival of acclimated crickets (compare to Toxopeus et al., 2019c), and we propose that this was caused by the temperature RNAi protocol itself, as 3 d at 22°C post-acclimation causes a decrease in freeze tolerance (Fig. 4).

**Figure 7.**
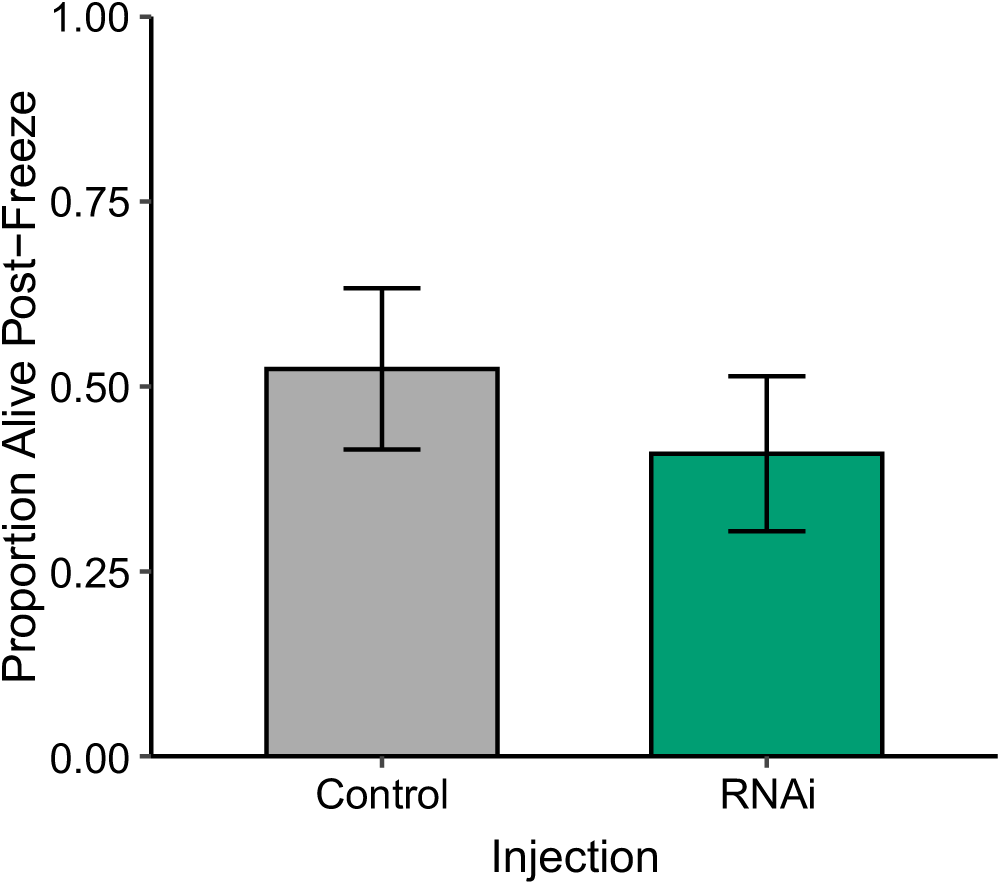
*Catalase* knockdown has no effect on freeze tolerance of acclimated crickets. Acclimated crickets were injected with dsRNA targeting *Catalase* (RNAi) or a control construct, and remained at a warm (c. 22°C) temperature for 3 days prior to testing freeze tolerance. The proportion of crickets that survived freezing was determined 48 h after exposing them to a 1.5 h freeze treatment at -8°C. Each treatment group included 21 – 22 crickets. Error bars represent the standard error of proportion. There was no effect of RNAi treatment on proportion survival based on a χ^2^ test (P < 0.05).

## Discussion

Here, we report the first use of RNAi in a freeze-tolerant insect to test whether a target gene is important for surviving freezing. Consistent with previous transcriptomic results (Toxopeus et al., 2019a), we showed that *Catalase* was upregulated at the mRNA level in fat body during fall-like acclimation, which likely supported the increased activity of the Catalase enzyme over the same time period, and justified the choice of *Catalase* as a target for RNAi (Lebenzon and Toxopeus, 2024). *Catalase* knockdown via RNAi was both tissue- and temperature-specific, and was most effective in fat body and midgut at warm (c. 22°C) temperatures. However, there was no difference in freeze tolerance (survival following a mild freeze treatment) in acclimated crickets that were injected with control dsRNA constructs (regular *Catalase* expression and activity) or *Catalase* dsRNA (*Catalase* knocked down). Therefore, we did not find support for our hypothesis that oxidative stress tolerance is important for freeze tolerance in *G. veletis*.

### Temperature- and time-dependence of RNAi efficacy

While many studies have shown that RNAi efficacy can vary over time, few have examined the effect of temperature on this biological process (Lebenzon and Toxopeus, 2024). Because *Catalase* knockdown was effective at warm (c. 22°C) but not moderately cool (15°C) temperatures, we conclude that RNAi is temperature-dependent in *G. veletis*. The limited efficacy of RNAi at low temperatures is similar to what has been observed in some insects (Fortier and Belote, 2000) and plants (Szittya et al., 2003), but differs from studies that show effective RNAi at low temperatures (Koštál and Tollarová-Borovanská, 2009; Romon et al., 2013; Vatanparast et al., 2022). A few hypotheses have been articulated about why RNAi may be inhibited at low temperatures (Lebenzon and Toxopeus, 2024). First, low temperatures decrease membrane fluidity, which could inhibit dsRNA transport into cells (Rode et al., 1997; Wolkers et al., 2003). Second, the RNAi process relies on a large suite of enzymes (Wilson and Doudna, 2013), some of which may function poorly at lower temperatures. Finally, RNAi relies on base pairing between nucleic acids (e.g., the guide RNA and the mRNA targeted for degradation; Wilson and Doudna, 2013), and nucleic acid base pairing is often temperature dependent. Our results hint that RNAi may be simply slowed down by the cold in *G. veletis*, as there was a trend toward decreased *Catalase* mRNA abundance 7 d post-injection at 15°C, suggesting that longer incubations with the dsRNA constructs could eventually lead to effective RNAi (e.g., 20 d; Des Marteaux et al., 2022; Ikeno et al., 2010). This is an important future direction, as even a short (3 d) period at warm temperatures post-acclimation was correlated with a loss of freeze tolerance in approximately 50% of crickets. The rapid deacclimation associated with brief warming (similar to Bennett and Lee, 1989; Williams et al., 2014) highlights a challenge associated with optimizing an RNAi protocol in a species that must be kept under conditions (cool temperatures) that are important for maintaining the phenotype of interest (freeze tolerance), but that inhibit RNAi efficacy.

### Effect of Catalase knockdown on freeze tolerance

While our RNAi protocol itself (e.g., 3 d at 22°C) may have affected freeze tolerance, the knock down of *Catalase* in acclimated crickets did not, which suggests there is minimal need for antioxidant protection during freezing and thawing in *G. veletis*. Studies that document oxidative damage in insects after freezing and thawing are rare, but Doelling et al. (2014) have shown that protein carbonylation and lipid peroxidation (markers of oxidative damage) increase after repeated freeze-thaw cycles. Thus, one future direction is to test whether oxidative damage occurs in *G. veletis* following differing intensities, durations, and frequencies of freezing events (see Schulte, 2014). It is also possible that low Catalase activity is sufficient to protect against freezing-associated oxidative damage, or that other antioxidants are more important for freeze tolerance in *G. veletis*. While *Catalase* was the only antioxidant gene upregulated in the fat body transcriptome of acclimated crickets (Toxopeus et al., 2019a), other antioxidants should be studied – especially in other tissues – to determine their possible contribution to freeze tolerance. Because freezing can damage mitochondria (e.g., Štětina et al., 2020), superoxide production from damaged mitochondria may be a more serious threat to survival than H_2_O_2_, and therefore Superoxide Dismutase may be a stronger candidate for facilitating freeze tolerance than Catalase.

### Tissue-specific expression and RNAi targeting Catalase

The tissue-specific expression and RNAi efficacy of genes has been recorded by others (Lebenzon and Toxopeus, 2024), and our results are consistent with much of the literature on this topic. In *G. veletis*, we hypothesize that some of the tissue-specific results were caused by differences in metabolism or other properties (e.g., other antioxidant defenses) in each tissue. Fat body tissue is lipid-rich and generally metabolically active (Arrese and Soulages, 2010). We speculate that the increased *Catalase* expression and activity during acclimation in fat body tissue of *G. veletis* is predominantly to mitigate H_2_O_2_ produced by lipid catabolism (Shields et al., 2021). We did not see elevated expression or activity of our target in midgut tissue, contrary to studies that show relatively high *Catalase* expression or activity in midguts of some insects (Felton and Duffey, 1991; Rajarapu et al., 2011), but consistent with relatively low expression in midguts of locusts (Zhang et al., 2016). It is possible that other antioxidants (e.g., glutathione; Barbehenn, 2003) are more important in this tissue in *G. veletis*, or that acclimation does not induce substantial H_2_O_2_-related oxidative stress in midgut. Malpighian tubules are metabolically active and express multiple detoxification-related genes (Nocelli et al., 2016). Our results also show that Catalase activity is higher in this tissue than the others we studied (fat body, midgut), similar to what is seen in locusts (Zhang et al., 2016). Interestingly, we see stable Catalase activity across acclimation in this tissue, even when *Catalase* mRNA abundance decreases in late acclimation. This is consistent with our finding that RNAi decreased transcript abundance in Malpighian tubules of unacclimated crickets, but enzyme activity remained unaffected. We won’t make any broad conclusions that Malpighian tubules are generally recalcitrant to RNAi while fat body and midgut are permissive to RNAi because our results may be particular to the target gene (*Catalase*) and tissue-specific mechanisms of regulating this gene.

### Conclusions

Our study shows that RNAi is possible in a freeze-tolerant insect, which will facilitate future studies examining the mechanisms underlying insect freeze tolerance. We also showed that RNAi was temperature- and tissue-specific, which will inform future RNAi protocols in *G. veletis*. While knockdown of *Catalase* did not affect survival of a mild freeze treatment, subsequent experiments could examine other antioxidants or use different freeze treatments to explore the relationship between freezing and oxidative stress.

## Acknowledgements

The authors would like to thank P. Brown, K. Lemay, L. McIntyre, M. Moore, M.L. van Oirschot and the StFX Animal Care Facility for help with rearing and/or processing crickets for our experiments, and N. Pichaud for advice on Catalase activity assays.

## Author Contributions

**SER**: conceptualization, methodology (RNAi optimization), investigation (RNAi enzyme assays), writing – original draft, writing – review and editing. **VEA**: investigation (RNAi qPCR), data curation, writing – review and editing. **RW**: conceptualization, methodology (enzyme assay optimization), investigation (acclimation enzyme assays), writing – original draft, writing – review and editing. **GEK**: conceptualization, methodology (qPCR optimization), investigation (acclimation qPCR), writing – original draft, writing – review and editing. **ALG**: investigation (RNAi qPCR), data curation, writing – review and editing. **JT**: conceptualization, data curation, funding acquisition, supervision, visualization, writing – original draft, writing – review and editing.

## Data and Code Availability Statement

All data and analysis code associated with this manuscript are available at https://github.com/jtoxopeus/cricket-catalase-rnai

